# Hawkes process modeling quantifies complicated firing behaviors of cortical neurons during sleep and wakefulness

**DOI:** 10.1101/2023.07.29.550297

**Authors:** Takeshi Kanda, Toshimitsu Aritake, Kaoru Ohyama, Kaspar E. Vogt, Yuichi Makino, Thomas J. McHugh, Hideitsu Hino, Shotara Akaho, Noboru Murata

**Affiliations:** Department of Neurophysiology, Nara Medical University, 840 Shijocho, Kashihara, Nara 634-8521, Japan; Institute for Integrative Sleep Medicine, University of Tsukuba, 1-1-1 Tennodai, Tsukuba, Ibaraki 305-8575, Japan; Hitotsubashi Institute for Advanced Study, Hitotsubashi University, 2-1 Naka, Kunitachi-shi, Tokyo 186-8601, Japan; National Institute for Materials Science, 1-1 Namiki, Tsukuba, Ibaraki 305-0044 Japan; Laboratory for Circuit and Behavioral Physiology, RIKEN Center for Brain Science, 2-1 Hirosawa, Wako, Saitama 351-0198, Japan; The Institute of Statistical Mathematics, 10-3 Midori-cho, Tachikawa, Tokyo 190-8562, Japan; National Institute of Advanced Industrial Science and Technology, 1-1-1 Umezono, Tsukuba, Ibaraki 305-8568, Japan; Waseda University, 3-4-1 Ookubo, Shinjuku, Tokyo 169-8555, Japan

## Abstract

Despite the importance of sleep to the cerebral cortex, how much sleep changes cortical neuronal firing remains unclear due to complicated firing behaviors. Here we quantified firing of cortical neurons using Hawkes process modeling that can model sequential random events exhibiting temporal clusters. “Intensity” is a parameter of Hawkes process that defines the probability of an event occurring. We defined the appearance of repetitive firing as the firing intensity corresponding to “intensity” in Hawkes process. Firing patterns were quantified by the magnitude of firing intensity, the time constant of firing intensity, and the background firing intensity. The higher the magnitude of firing intensity, the higher the likelihood that the spike will continue. The larger the time constant of firing intensity, the longer the repetitive firing lasts. The higher the background firing intensity, the more likely neurons fire randomly. The magnitude of firing intensity was inversely proportional to the time constant of firing intensity, and non-REM sleep increased the magnitude of firing intensity and decreased the time constant of firing intensity. The background firing intensity was not affected by the sleep/wake state. Our findings suggest that the cortex is organized such that neurons with a higher probability of repetitive firing have shorter repetitive firing periods. In addition, our results suggest that repetitive firing is ordered to become high frequency and short term during non-REM sleep, while unregulated components of firing are independent of the sleep/wake state in the cortex. Hawkes process modeling of firing will reveal novel properties of the brain.

## 1 INTRODUCTION

The sleep/wake state modulates firing behavior of neurons in the cerebral cortex, making a difference in cortical functions during sleep and wakefulness (Levenstein et al., 2017; Steriade and Timofeev, 2003). The manner in which neuronal firing fluctuates is critical for signal processing in the brain, such as determining the direction and duration of synaptic plasticity (Froemke and Dan, 2002; Sjöström et al., 2001). Sleep-induced changes in cortical neuronal firing are, however, not well quantified due to the complicated behavior. Non-rapid-eyemovement sleep (non-REM) sleep, a sleep state characterized by slow waves on the electroencephalogram (EEG), elicits large oscillations called slow-wave activity (SWA) in the cortical local field potentials (LFP) and rapid transition of depolarization and hyperpolarization called the UP/DOWN states in membrane potentials of individual cortical neurons (Destexhe et al., 1999; Steriade et al., 2001; Timofeev et al., 2001). In contrast to these characteristic intracellular and extracellular oscillations, individual cortical neuronal firing does not exhibit constant oscillatory behavior during non-REM sleep, which is due to highly diverse firing responses to depolarization and hyperpolarization in cortical neurons (Connors and Gutnick, 1990). To understand the precise behavior of the sleeping cerebral cortex at the individual cellular level, in this study, we quantify the complicated firing of cortical neurons during wake and non-REM sleep using Hawkes process, a statistical model used for modeling non-equidistant event time series.

Hawkes process, a temporal point process proposed by Hawkes (Hawkes, 1971b,a), is used to model phenomena in which sequential events are concentrated in a short period, such as earthquakes (Adamopoulos, 1976; Ogata, 1988, 1998), financial markets (Hawkes, 2018), e-mailing (Fox et al., 2016), viral diffusion on social media (Kobayashi and Lambiotte, 2016), spread of infections (Chiang et al., 2022; Meyer et al., 2012; Koyama et al., 2021), urban crimes (Mohler et al., 2011), and neuronal network dynamics (Chornoboy et al., 1988; Pernice et al., 2011). In Hawkes process, “intensity”, a parameter that defines the probability of an event occurring, is self-exciting, where the intensity temporarily increases with each occurrence of an event, causing events to occur in clusters. In this report, we defined the appearance of repetitive firing as the firing intensity corresponding to “intensity” in Hawkes process. Three parameters, the magnitude of firing intensity, the time constant of firing intensity, and the background firing intensity, were used to quantify firing behavior of individual cortical neurons.

## 2 METHODS

### 2.1 Experiments

#### 2.1.1 Animals

All experimental procedures and protocols were approved by the Institutional Animal Care and Use Committee of the University of Tsukuba. Mice were fed ad libitum and maintained in a temperature- (22^*°*^C) and humidity-controlled room under a strict 12-h light-dark cycle (lights on 9:00 AM to 9:00 PM).

#### 2.1.2 Electrophysiology

The tetrode recordings in freely-moving mice were previously reported (Ohyama et al., 2020). Briefly, multi-unit activity was recorded with chronically implanted tetrode electrodes in the primary motor and somatosensory cortices of C57BL/6 mice during sleep and wakefulness. Electroencephalography (EEG) data were recorded using two screw electrodes on the skull. Electromyography (EMG) data were acquired with two stainless steel wires implanted into the neck muscle. All data were acquired with a multichannel amplifier (Digital Lynx SX, Neuralynx). Spike data were band-pass filtered between 0.6 and 6 Hz, and digitized at 32 kHz. EEG and EMG data were sampled at 250 Hz through a 0.1 Hz high-pass filter for EEG and a 10 Hz high-pass filter for EMG. Single units were isolated by manual cluster cutting with Spikesort3D (Neuralynx). EEG and EMG data were divided into 4-s time windows, and sleep/wake states were scored according to the following criteria: desynchronized EEG and high-amplitude EMG during wake, high-amplitude slow-wave EEG (1–4 Hz) and lower-amplitude EMG during non-REM sleep, and theta-frequency EEG (7–9 Hz) and much lower-amplitude EMG during REM sleep. Because of the low frequency of REM sleep in this study, data during REM sleep were excluded from the following analyses.

### 2.2 Statistical modeling

#### 2.2.1 Temporal Point Process

A temporal point process is a stochastic model for a time series of discrete events, where the events are treated as a set of non-equidistant timestamps 𝒯 = {*t*_*i*_} of the event occurrence. In this work, we model the above electrophysiology data as the temporal point process, where the event is firing of a neuron. This process is characterized by the conditional firing intensity function *λ*(*u*|ℌ_*u*_), where H_*u*_ is the set of firing timestamps before time *u*. Then, the probability of firing in some small interval [*u, u* + *du*] is defined as

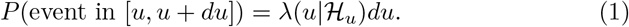

Namely, the probability of neuronal firing depends on the firing intensity function that includes influence from past firing.

When the intensity function is a constant function, which does not depend on the past firings, the temporal point process is called a Poisson process. Similarly, when the intensity function only depends on the current timestamp *u*, the temporal point process is called an inhomogeneous Poisson process. Henceforth, we shall write *λ*(*u*|ℌ_*u*_) by *λ*(*u*) as a shortened form when there is no risk of confusion.

Given a timestamp of the last firing *s*, let the cumulative distribution function of firing at time *u* ϵ [*s*, ∞) be *F* (*u*|ℌ_*s*_) and its density function be *f* (*u*|ℌ_*s*_). We remark that in this case ℌ_*s*_ is identical to H_*u*_, because no firing is observed in [*s, u*). Then the firing intensity function can be written as

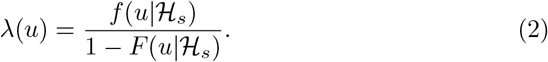

Integrating both sides of the above equation from *t*_*k*_ to *t* yields

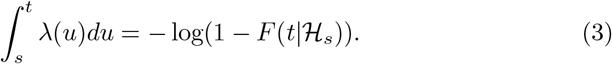

Therefore,

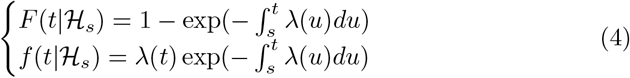

Then, the likelihood of firings 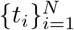, occurring in [0, *T*], can be calculated using chain-rule as follows, where *N* represents the total number of events occurring in [0, *T*].

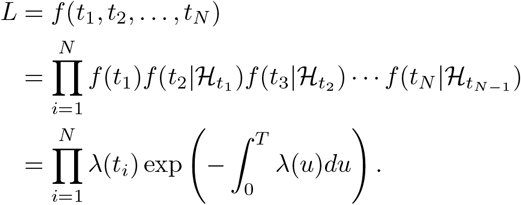

Then, the log-likelihood of the firings can be written as

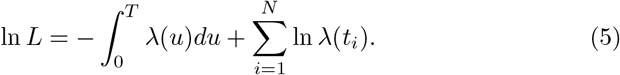

#### 2.2.2 Hawkes Process

In this report, Hawkes process, a self-exciting temporal point process, was used to model short-term burst firing. In Hawkes process, the following firing intensity function, which depends on the timestamps of past firings, is used.

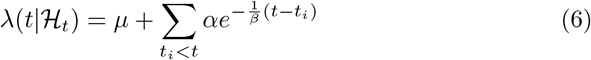

where *α* is the magnitude of firing intensity and *β* is the time constant of the firing intensity. Namely, the intensity value increases by *α* when firing and excites successive firings. Then, the increment of the intensity halves after *β* ln 2 time, and converges to 0 when *t* →+∞ unless no new firing occurs. The background firing intensity *μ*is a constant baseline firing intensity.

The log-likelihood of Hawkes process for firings 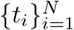 in [0, *T*] is given by Eq. (5). More concretely, the log-likelihood can be written as

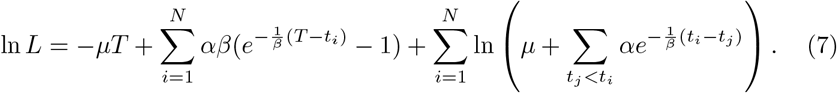

This log-likelihood can be differentiable with respect to *α, β*, and *μ*. Therefore, given a set of timestamps of firings 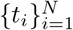 in [0, *T*], maximum likelihood estimation (MLE) of these parameters is possible, and recursive method to compute the gradient is provided by Ozaki (1979), and the asymptotic properties of maximum likelihood estimation are also analyzed (Ogata, 1978). However, the MLE tends to be numerically unstable due to the log term in Eq. (7), and *β* can take an abnormally large value. This means that the intensity of the Hawkes process does not decrease after a firing and the firing of infinitely many neurons is triggered. To avoid such solutions, estimation by EM algorithm is proposed (Veen and Schoenberg, 2008; Halpin and De Boeck, 2013). Let *α*^(*k*)^, *β*^(*k*)^, and *μ*^(*k*)^ be the estimated parameter after the *k*th iteration. In EM algorithm the log term in Eq. (7) is bounded below by

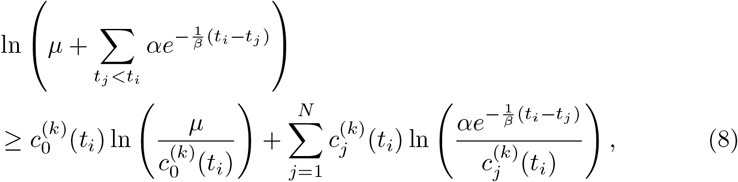

where

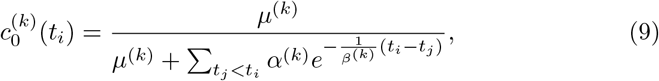

and,

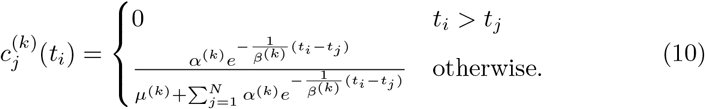

Here, the equality in Eq. (8) holds when *α* = *α*^(*k*)^, *β* = *β*^(*k*)^, and *μ*= *μ*^(*k*)^.

Then the log-likelihood in Eq. (7) is lower-bounded as

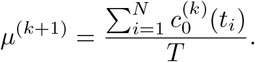

The parameters are updated by maximizing the lower-bound 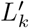. By setting the gradients of 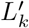 with respect to *μ* to zero, we obtain

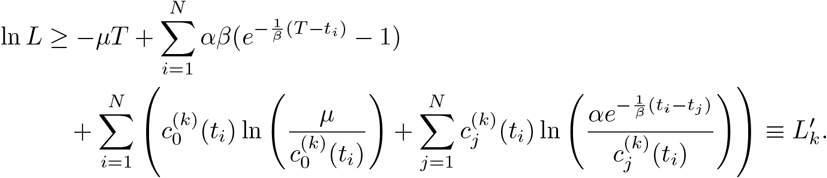

Similarly, by setting the gradients of 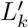 with respect to *α* and *β* to zeros, the following equations hold for the solution that maximizes the 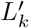.

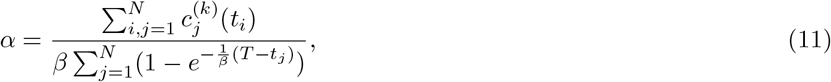

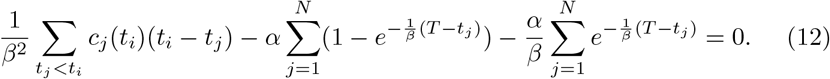

Substituting Eq. (11) into Eq. (12) yields the equation of *β*. We obtain *β*^(*k*+1)^ by solving the obtained equation using root-finding algorithms such as Newton’s Method, and *α* ^(*k*+1)^is obtained by substituting *β*^(*k*+1)^ into *β* in Eq. (11).

Then, we lower-bounds the log-term in Eq. (7) by Eq. (8) using 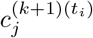 calculated by Eqs. (9) and (10), and compute the solution that maximizes the lower-bound 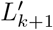. These steps are repeated until ‖***θ***^(*k*)^ − ***θ***^(*k*+1)^ ‖_2_ *< ϵ* is met for a sufficiently small number *ϵ*, where ***θ***^(*k*)^ = [*α*^(*k*)^, *β*^(*k*)^, *μ*^(*k*)^]^*T*^.

### 2.3 Data analysis

#### 2.3.1 Application to the electrophysiology data

Hawkes process assumes that the parameter ***θ*** = [*α, β, μ*]^*T*^ is invariant over the time interval [0, *T*], and only the intensity function varies over time depending on the event occurrence.

On the other hand, we assume that the parameter distribution changes depending on the sleep state *s*_*t*_ of time *t*. To handle the different parameters over time for the Hawkes process, we use sliding windows and estimate the parameter ***θ***_*t*_ for the events in each sliding window as in Fig. 2 B.

We chose *T* = 20-s as the length of the sliding window. The window was slid every 4-s. Continuous 20 s wake or non-REM periods were used for Hawkes process modeling. If there were less than 10 spikes in the 20-s time window, the firing for that time was not modeled by Hawkes process because the estimated parameters are unreliable. Hawkes process was applied to neurons that exhibited at least five each of wake and non-REM periods that could be modeled by Hawkes process (≥10 spikes/20-s). The time window and data utilization criteria were used not only for Hawkes process but also for analyzing firing rate and interspike intervals.

#### 2.3.2 Statistical tests

For the firing rate and the coefficient of variation of interspike intervals (CV of ISI), in this study, the mean of each parameter was used as the representative value of individual neurons. For the firing intensity parameters, in this work, the representative value of individual neurons was the median of each parameter, because the distribution was heavy-tailed. Wilcoxon signed-rank test was used to compare two paired groups (Fig. 1 B and C, Fig. 3 A). The significance level was set at 0.05. In Fig. 3 B and subsequent figures, independence between two groups was tested by Hilbert-Schmidt independence criterion (HSIC) (Gretton et al., 2005), indicating that all pairs were not independent (*p* < 3.00 × 10^*−*3^). The relationship between the two groups was evaluated using the maximal information coefficient (MIC) (Reshef et al., 2011). The values in this report were presented as mean ± standard deviation.

**Figure 1:**
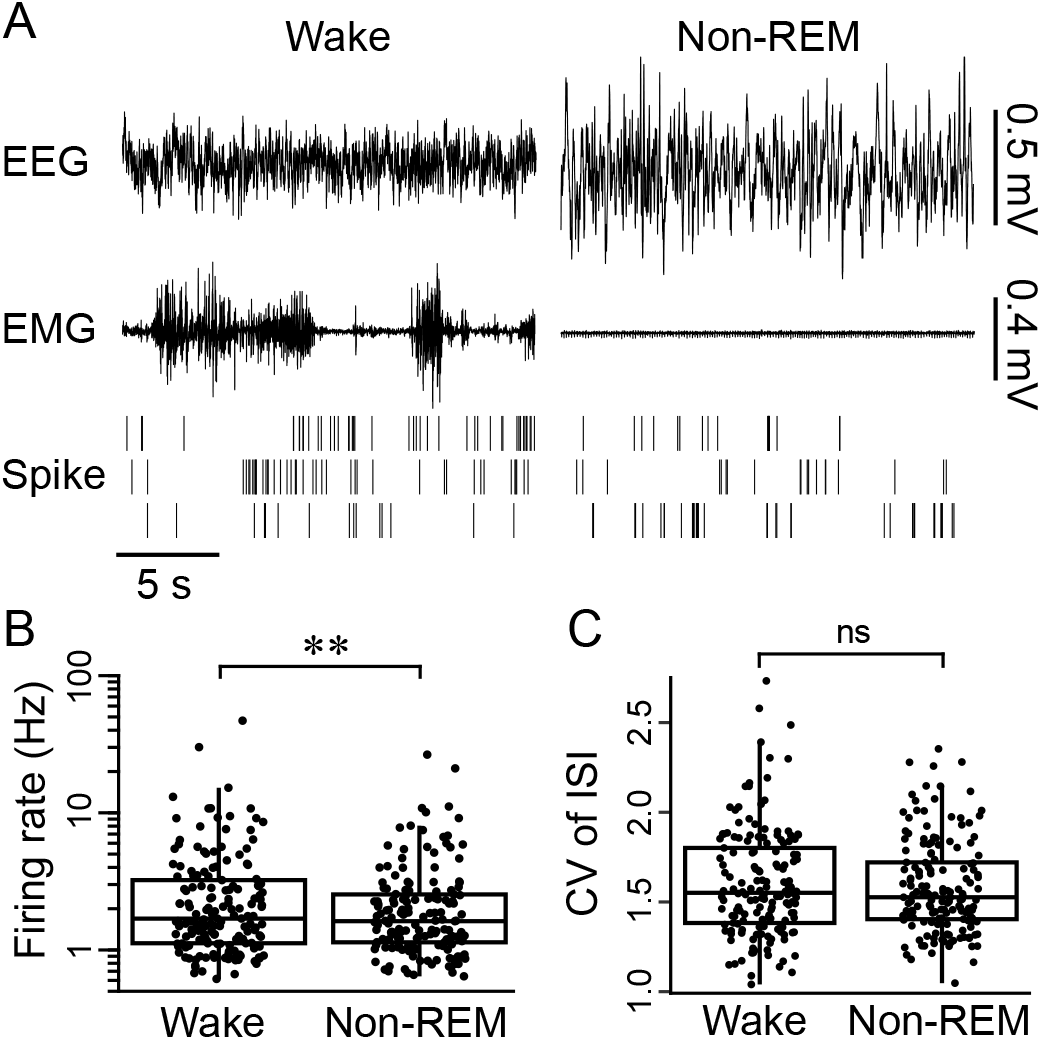
Basic properties of cortical neuronal firing during wake and non-REM sleep. (A) Multi-unit recording from layer 5 of the primary motor cortex during wake and non-REM sleep. (upper and middle) Simultaneously recorded EEG and EMG. (lower) The raster plots of unitary activity. Each row indicates spikesorted units from tetrode electrode data. (B and C) Box plots with mean values of individual cortical neurons during wake and non-REM sleep. Dots indicate the mean values. (B) Firing rates with a logarithmic scale. ∗∗, *p* < 0.01. (C) Coefficient of variation of inter-spike intervals (CV of ISI). ns, *p* > 0.05.

**Figure 2:**
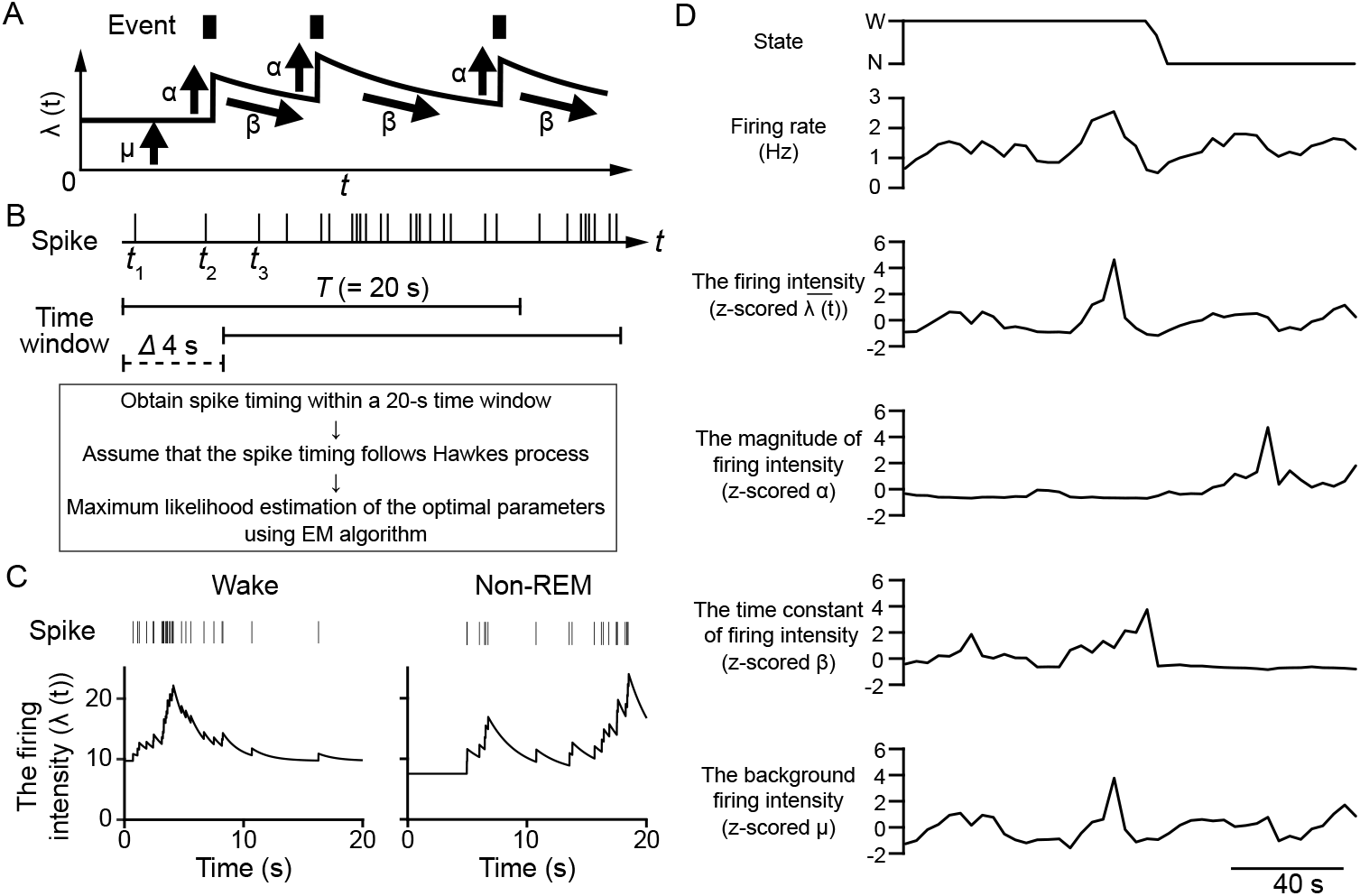
Application of Hawkes process modeling to firing pattern analysis. (A) A schematic showing of Hawkes process. Each parameter corresponds to Eq. (6). (B) A diagram of the analysis in this study. (upper) Spike timing. *t*_*i*_ indicates the time of the *i*th spike occurrence. (middle) Time window for 20-s every 4-s. (lower) The pipeline for analysis of firing patterns using Hawkes process. EM algorithm is used to estimate the firing intensity parameters. (C) Representative traces of the firing intensity (*λ*(*t*)) in a 20-s time window during wake and non-REM sleep. Left and right data are taken from the same neuron. (upper) Spike timing. (lower left) *λ*(*t*) of *α*=1.1, *β*=0.7, and *μ*=0.3. (lower right) *λ*(*t*) of *α*=2.0, *β*=0.4, and *μ*=0.4. (D) Representative traces of the firing rates and intensity parameters during the transition from wake to non-REM sleep. The calculation is based on 20-s time windows that move in 4-s steps. The values, except for firing rate, were z-scored against each value for 20-s. The values of 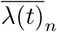 in this graph are the mean of *λ*(*t*) every time window. States are shown on the top: W=wake, N=non-REM. The intermediate state is a state in which there are both wake and non-REM in a 20-s time window.

**Figure 3:**
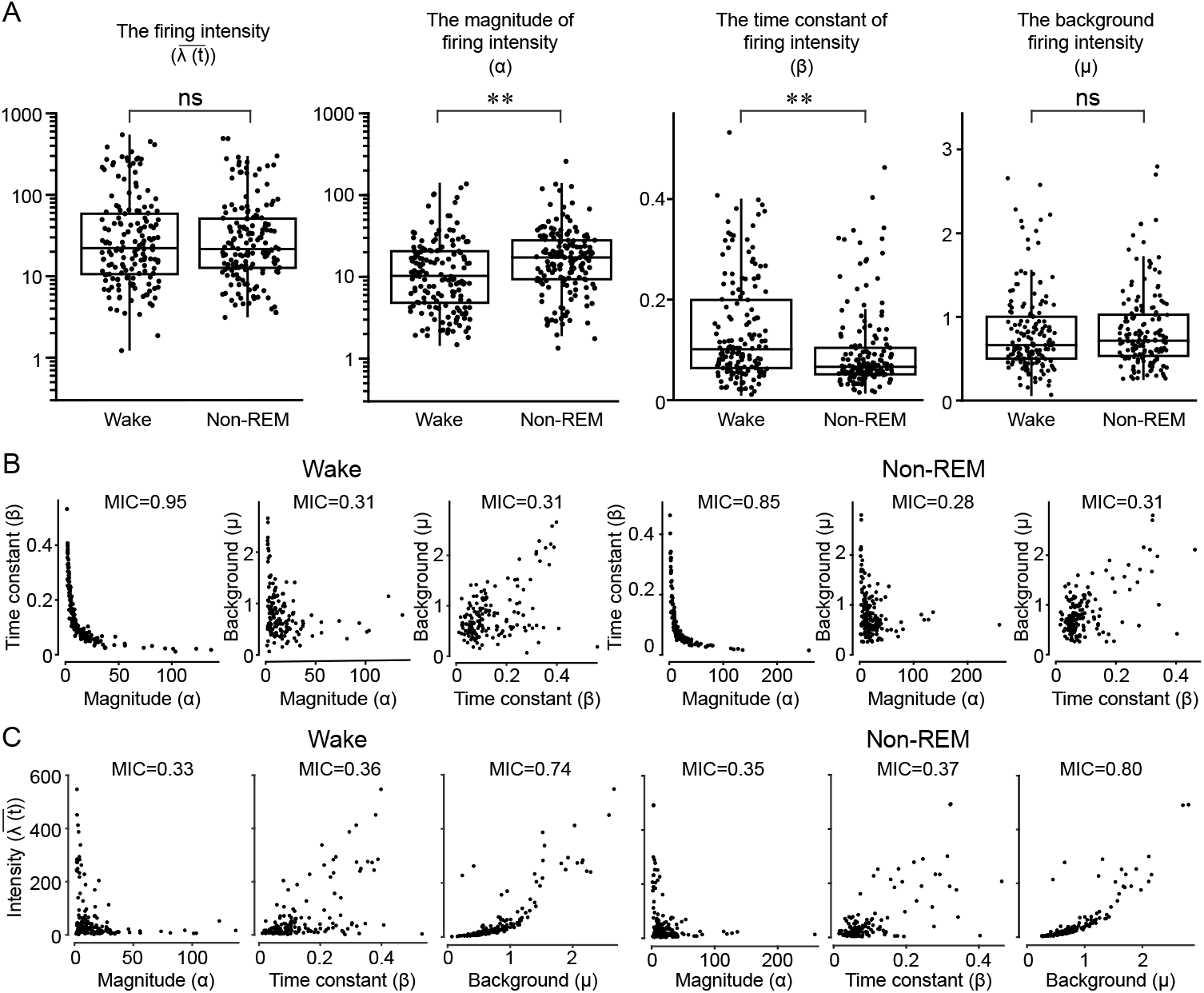
Quantifying firing behaviors of cortical neurons during wake and non-REM sleep with the firing intensity parameters. (A) Box plots with median values of the firing intensity parameters in individual cortical neurons during wake and non-REM sleep. Dots indicate the median values of individual neurons. The two left panels are logarithmic scales. ∗∗, p < 0.01. ns, p > 0.05. (B and C) Scatter plots for each firing intensity parameter in wake (left) and non-REM sleep (right). The maximal information coefficient (MIC) is shown at the top of each panel.

## 3 Results

### 3.1 Basic characterization of firing behavior of cortical neurons during wake and non-REM sleep

Using existing methods, we first characterized firing of 161 neurons in the somatosensory and motor cortices acquired from 6 mice with tetrode recordings. Their firing rate was slightly but significantly higher during wake than during non-REM sleep (3.17 ± 4.83 Hz in wake, 2.55 ± 3.12 Hz in non-REM, *p* = 5.82 × 10^*−*6^, Fig. 1 B). A few little changes in averaged individual cortical activity between wake and non-REM sleep are observed in a number of previous reports (Evarts, 1964; Vyazovskiy et al., 2009; Ohyama et al., 2020; Miyazaki et al., 2020; Seibt et al., 2017; Hengen et al., 2016). Non-REM sleep exerts the UP/DOWN oscillations in the membrane potentials of cortical neurons, which could change their firing patterns. There was, however, no significant difference in the coefficient of variation of interspike intervals (CV of ISI), an evaluation method for firing patterns, between wake and non-REM sleep (1.60 ±0.30 in wake, 1.58 ±0.26 in non-REM, *p* = 2.57 × 10^*−*1^, Fig. 1C), suggesting that changes in firing patterns of cortical neurons during non-REM sleep are not well reflected in the CV of ISI.

### 3.2 Hawkes process modeling of cortical neuronal firing

To further understand the sleep/wake state-dependent behaviors of cortical neurons, we next explored the individual firing patterns using Hawkes process. Hawkes process is a useful statistical model to investigate sequential events, which was applied in this study to quantify repetitive firing of cortical neurons. Figure 2 A shows a schematic of Hawkes process. If occurrences of events follow Hawkes process, event timings are characterized by Eq. (6) (see METHODS). The larger *λ*(*t*), the more likely an event is to occur, and a baseline value *μ*of *λ*(*t*) causes uniformly random events as a Poisson process. When an event occurs, *λ*(*t*) increases by *α*which triggers successive occurrences of events. The increment decays with *t* at the rate of *β*.

We defined the appearance of repetitive firing as the firing intensity *λ*(*t*), and behaviors of cortical neurons were quantified by the three parameters (herein referred to as the firing intensity parameters): the magnitude of firing intensity *α*, the time constant of firing intensity *β*, and the background firing intensity *μ* for 20-s time windows (Fig. 2 B). The magnitude of firing intensity denotes the influence of past firing on the following firing occurrence; the higher the magnitude of firing intensity, the more likely it is that repetitive firing will occur. The time constant of firing intensity represents the time constant for the decay of the magnitude of firing intensity once increased; the lower the time constant of firing intensity, the shorter the duration of repetitive firing is. The background firing intensity indicates the constant firing intensity that does not depend on the occurrence of firing; the higher the background firing intensity is, the higher the firing intensity constantly is. The firing intensity parameters were estimated by EM algorithm. Figure 2 C shows traces of the firing intensity determined by the firing intensity parameters estimated from spike timing in the 20-s time window. In the time window of Figure 2 C, the firing intensity exhibited a stepwise increase and exponential decay for each spike as per Figure 2 A, which was characterized by the slower decay during wake (*β*, 0.7 in wake; 0.4 in non-REM) and the higher increase during non-REM sleep (*α*, 1.1 in wake; 2.0 in non-REM). Figure 2 D shows traces of the firing rates and intensity parameters during the transition from wake to non-REM sleep. The firing intensity parameters fluctuated over time and depended on the sleep/wake states: in the case of the neuron in Fig. 2 D, the magnitude of firing intensity *α* increased gradually and the time constant of firing intensity *β* was suppressed during non-REM sleep. During the transient increase in the firing rate, similar changes were observed in the background firing intensity *μ*(Fig. 2D).

### 3.3 Quantification of firing patterns of cortical neurons with the firing intensity parameters

The firing intensity parameters were compared collectively between wake and non-REM sleep (Fig. 3 A). The magnitude of firing intensity was higher in non-REM than in wake (*α*, 17.0 ± 23.6 in wake, 21.5 ± 28.8 in non-REM, *p* = 2.60×10^*−*7^). The time constant of firing intensity was smaller in non-REM than in wake (*β*, 0.14 ± 0.10 in wake, 0.10 ± 0.08 in non-REM, *p* = 1.10 × 10^*−*12^). These results indicate that cortical neurons exhibit a higher probability to fire repetitively for shorter periods in non-REM sleep than in wake. The difference in the background firing intensity did not reach statistical significance (*μ*, 0.81± 0.84 in wake, 0.49 ±0.46 in non-REM, *p* = 6.18 × 10^*−*2^). Since the background firing intensity *μ*is, in other words, the intensity of a Poisson process underlying firing, an ability to fire randomly in cortical neurons may not depend on the sleep/wake states. The mean firing intensity 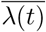, the mean value of firing intensity *λ*(*t*) for the time window in [*t, t* + 20], is by nature dependent on the background firing intensity *μ*, thus, as expected, the mean firing intensity also did not change during wake and non-REM sleep 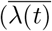, 62.5 ±52.9 in wake, 97.2 ±81.1 in non-REM, *p* = 1.83 × 10^*−*1^). These results indicate that Hawkes process modeling can quantify the sleep/wake-state-dependent changes in firing patterns of cortical neurons that are not reflected by CV of ISI (Fig. 1 C).

The firing intensity parameters are independent variables in Hawkes process modeling, but the interrelationships in neuronal firing need to be examined for understanding the properties of modeled firing patterns. We next investigated the relationship between the firing intensity parameters. Since a nonlinear relationship was observed in some cases (Fig. 3 B), the relationship was evaluated by the maximal information coefficient (MIC) that can detect both linear and nonlinear correlations between two variables (Gretton et al., 2005). The magnitude of firing intensity *α* was inversely proportional to the time constant of firing intensity *β* in both wake and non-REM sleep (MIC = 0.95 in wake, MIC = 0.85 in non-REM), while no close correlation was found between the other firing intensity parameters (MIC ≤ 0.31, Fig. 3 B), suggesting that, in the cerebral cortex, neurons with high probability of repetitive firing are less likely to sustain repetitive firing over the long term. We next examined which and how much firing intensity parameters influence the mean firing intensity 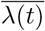. As expected, a strong correlation between the mean firing intensity 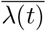 and the background intensity *μ*was observed in both wake and non-REM sleep (MIC = 0.74 in wake, MIC = 0.80 in non-REM), whereas the other firing intensity parameters showed only slight correlation with the mean firing intensity 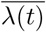 (MIC≤ 0.37, Fig. 3 C), confirming that the background firing intensity *μ*is the factor with the greatest impact on the mean firing intensity 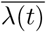.

### 3.4 The relationship between the firing intensity parameters across the sleep/wake cycle

We finally examined the relationship between the firing intensity parameters in the different states. In this section, *x*_*w*_ and *y*_*n*_ indicate the parameter *x* in wake and *y* in non-REM, respectively. In the comparison of the same parameters between wake and non-REM, a relatively strong correlation was found for the mean firing intensity 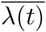 and the background firing intensity *μ*(MIC = 0.61 between 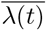 and 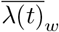, MIC = 0.57 between *μ*_*w*_ and *μ*_*n*_, in Fig. 4). Although there were also positive correlations between the mean firing intensity 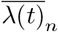 and the background firing intensity *μ* across different states (MIC = 0.57 between 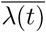 and *μ*_*n*_, MIC = 0.51 between *μ*_*w*_ and 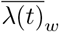, in Fig. 4), the correlations were weaker than those between the same states (Fig. 3 C). The relatively weak correlations were observed in the magnitude of firing intensity *α* or the time constant of firing intensity *β* between the two different states (MIC = 0.41 between *α*_*w*_ and *α*_*n*_, MIC = 0.44 between *β*_*w*_ and *β*_*n*_, in Fig. 4). The negative correlation between the magnitude of firing intensity and the time constant of firing intensity was strong during the same states (Fig. 3 C) but weakened across the different states (MIC = 0.41 between *α*_*w*_ and *β*_*n*_, MIC = 0.41 between *β*_*w*_ and *α*_*n*_, Fig. 4), suggesting that the individual characteristics of the rise and fall of the firing intensity following a single spike may not be conserved across the sleep/wake cycle. The other relationships between the firing intensity parameters during the different states were not strong (MIC *<* 0.40, Fig. 4).

**Figure 4:**
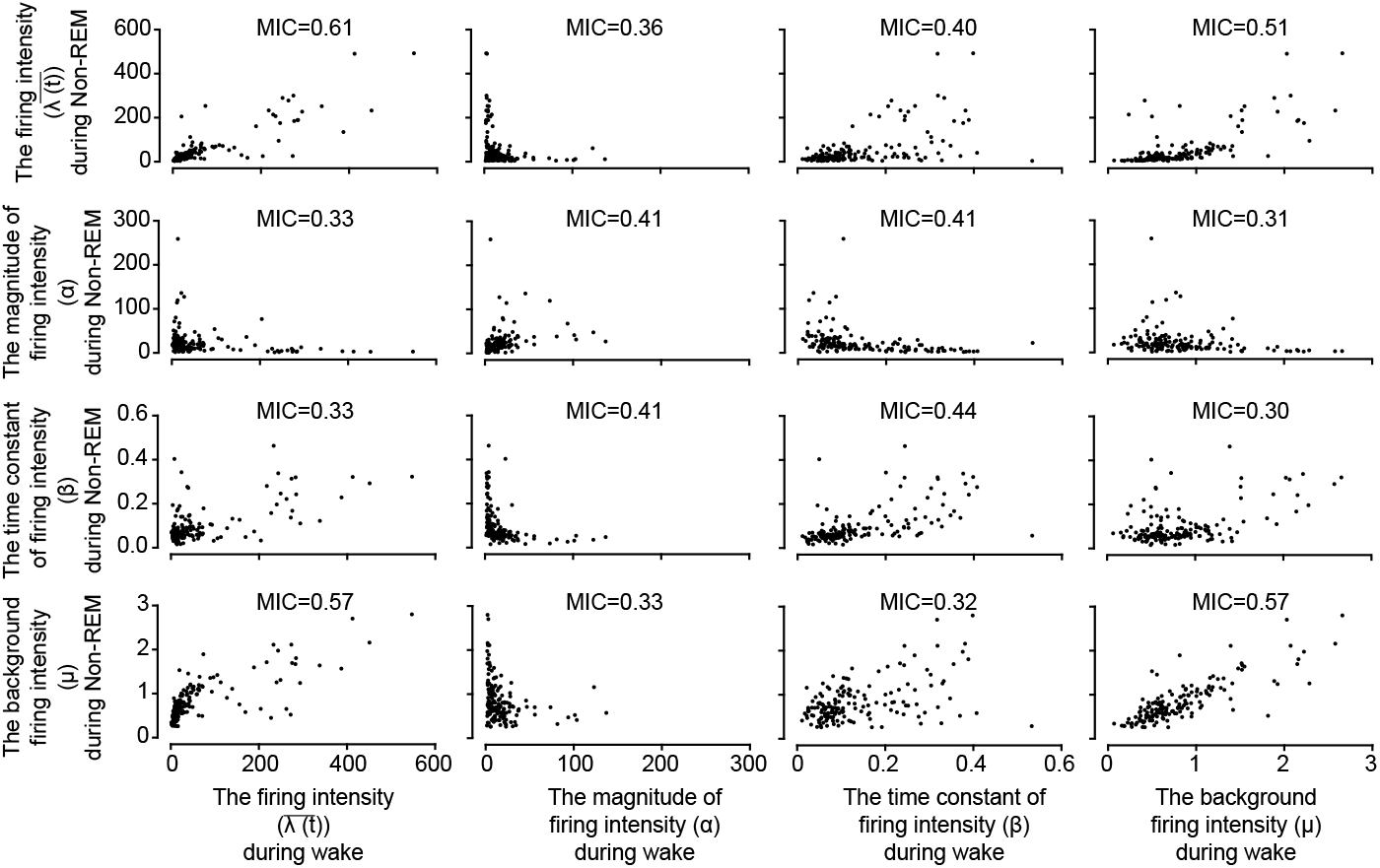
Relationship between the firing intensity parameters in wake and non-REM sleep. Scatter plots for the firing intensity parameters during wake (the horizontal x-axis) and non-REM sleep (the vertical y-axis). Dots indicate the median values of individual neurons. MIC is shown at the top of each panel.

## 4 DISCUSSION

### 4.1 Hawkes process modeling for complicated firing pattern analysis

Wake and non-REM sleep are distinct states from each other in the brain, which is evident from population dynamics in cortical neurons such as the EEG, the LFP, and the cell assembly (Loomis et al., 1935; Destexhe et al., 1999; Hengen et al., 2016; Seibt et al., 2017; Miyazaki et al., 2020; Nagayama et al., 2022). Despite the UP/DOWN oscillations in the membrane potentials during non-REM sleep (Destexhe et al., 1999; Steriade et al., 2001; Timofeev et al., 2001), it was not clear how much firing patterns of individual neurons, a fundamental process underlying brain functions, fluctuate across the sleep/wake cycle. Sleep is a spontaneous and intermittent physiological phenomenon, and the onset of sleep/wake states cannot be determined in milliseconds even using the EEG or the LFP. Thus, it is a bit problematic to analyze the sleep/wake state-related firing patterns with event-aligned methods such as peristimulus time histogram (PSTH) and spike-triggered average (STA). In some cases, the variance and the distribution of interspike intervals (ISI) differ in the cerebral cortex during sleep and wakefulness (Evarts, 1964; Hengen et al., 2016), but these do not always reflect firing patterns. Using Hawkes process modeling, we achieved to quantify the sleep/wake state-dependent changes in cortical neuronal firing and discovered the novel firing properties: a low-frequency and long-term repetitive firing during wake, a high-frequency and short-term repetitive firing during non-REM sleep, and a constant random firing independent of the sleep/wake state (Fig. 2 and 3). Our findings demonstrate that Hawkes process modeling is also useful for the analysis of complicated spiking in individual neurons and that wake and non-REM sleep make a distinct difference in individual firing patterns as well as known population dynamics and membrane potential fluctuations in the cerebral cortex. The SWA of 1–4 Hz emerges in the cortical LFP during non-REM sleep, which is often used to characterize cortical spikes such as the phase-locked firing and the correlation between firing rate and SWA power (Watson et al., 2016; McKillop et al., 2018; Xu et al., 2019; Ohyama et al., 2020; Thomas et al., 2020). Hawkes process modeling allowed the characterization of firing patterns of individual neurons without the use of the LFP, which is a great advantage that can be applied to analyze neuronal firing in the brain areas where characteristic LFP does not occur.

### 4.2 Neurophysiological insights into novel firing behaviors in the sleeping cortex

The magnitude of firing intensity *α* was inversely proportional to the time constant of firing intensity *β* in cortical neurons (Fig. 3 B), which is rational from the perspective of neuronal energy consumption and excitotoxicity. In the mammalian brain, neurons should not fire burst of action potentials for prolonged periods because excessive repetitive firing and synaptic transmission consume large amounts of adenosine triphosphate (ATP) to restore ionic composition (Harris et al., 2012) and, in the worst case, induces neuronal death through Ca^2+^ overload (Choi, 1992). The inversely proportional relationship between the magnitude and the time constant of firing intensity was observed in both wake and non-REM sleep (Fig. 3 B), allowing neurons to maintain self-protection against hyperexcitation regardless of the sleep/wake state that results in substantial changes. Non-REM sleep increased the magnitude of firing intensity and decreased the time constant of firing intensity (Fig. 3 A), which could be due to the UP/DOWN oscillations of the membrane potentials. Repetitive action potentials may occur in the intermittent UP state during non-REM sleep rather than in the sustained UP state during wake, because less repolarization lowers the driving force behind action potential generation. In fact, burst firing (ISI *<* 15 ms) of cortical neurons increases during non-REM sleep (Watson et al., 2016; Ohyama et al., 2020). Although the periodic DOWN state might cease firing of cortical neuron for a short period, it was never detectable from extracellular single unit data alone. Highly unstable ISI of cortical neurons, caused by the complicated spontaneous firing, is thought to be a factor that a short-term cessation of firing has been undetectable from ISI. The OFF period is a period of cessation of firing of neuronal populations, not individual neurons in the cerebral cortex, which emerges during non-REM sleep (Vyazovskiy et al., 2009; Watson et al., 2016). Since the UP/DOWN state is synchronized between neurons (Volgushev et al., 2006; Chauvette et al., 2010), the OFF period might correspond to the DOWN state. However, the OFF periods have been determined in a qualitative way from population firing or defined from the LFP (Vyazovskiy et al., 2009; Watson et al., 2016; McKillop et al., 2018). Our approach discovered from single unit data the repetitive firing does not last longer during non-REM sleep than during wake (Fig. 3 A), suggesting the existence of a cessation period of firing at the individual neuron level during non-REM sleep that cannot be detected by existing methods. This indicates high inference capabilities of Hawkes process modeling in firing pattern analysis. The background firing intensity *μ* did not depend on the sleep/wake state (Fig. 3 A), suggesting that changes in cortical neuronal firing elicited by the sleep/wake state are not due to an increase or decrease in Poisson-like fluctuations in membrane potentials and synaptic inputs. Hawkes process modeling is able to extract novel physiological characteristics of neurons even from single unit data only, which is applicable to firing analysis of neurons all over the brain under various conditions.

In this study, we focused on firing behavior of individual neurons without considering network structures. Presently, it is possible to record spikes of more than 1,000 neurons simultaneously (Steinmetz et al., 2021).

Hawkes process has also been extended to handle multivariate event series, allowing for the modeling of mutual influences among each event series. Therefore, it is possible to apply Hawkes process to the simultaneous multi-unit recordings and consider the neuronal network structures.

The application of machine learning or statistical methods including Hawkes process to large-scale spiking data or neuronal network structures will reveal undiscovered properties of the brain.

## 5 ACKNOWLEDGMENTS

Part of this study was carried out under the ISM Cooperative Research Program (2023-ISMCRP-1019). KO was supported by a Restart Postdoctoral Fellowship (RPD, 17J40129) from Japan Society for the Promotion of Science (JSPS).

## 6 DATA AVAILABILITY

Data will be made available upon reasonable request.

## 9 AUTHOR CONTRIBUTIONS

T.K., T.A., K.O., H.H., S.A., and N.M. conceptualized and designed the study; T.K., T.A., K.O., and K.E.V. performed experiments and analyzed data; Y.M. and T.J.M. developed experimental methodology; T.A., H.H., S.A., and N.M. performed modeling data; all the authors interpreted the results of experiments; T.K. and T.A. prepared the figures and wrote the first draft of the manuscript; all the authors edited and revised the manuscript and approved the final manuscript.

